# Attentional inhibition ability predicts neural representation during challenging auditory streaming

**DOI:** 10.1101/2022.09.29.510226

**Authors:** Joan Belo, Maureen Clerc, Daniele Schön

**Affiliations:** Centre Inria d’Université Côte d’Azur, Sophia Antipolis, France; Université Côte d’Azur, Nice, France; Institut de Neurosciences des Systèmes, Marseille, France; Institute for Language, Communication, and the Brain, Aix-en-Provence, France

## Abstract

Focusing on a single source within a complex auditory scene is challenging. M/EEG-based auditory attention detection allows to detect which stream, within a set of multiple concurrent streams, an individual is attending to. The high inter-individual variability in the AAD performance is most often attributed to physiological factors and signal to noise ratio of neural data. Here we address the hypothesis that cognitive factors and in particular sustained attention, WM and attentional inhibition, may also partly explain the variability in AAD performance, because they support the cognitive processes required when listening to complex auditory scenes. Here, we chose a particularly challenging auditory scene, by presenting dichotically polyphonic classical piano excerpts lasting one minute each. Two different excerpts were presented simultaneously in each ear. Forty-one participants, with different degrees of musical expertise, listened to these complex auditory scenes focussing on one ear while we recorded the EEG. Participants also completed several tasks assessing executive functions. As expected, attended stimuli were better decoded than unattended stimuli. Importantly, attentional inhibition ability did explain around 10% of the reconstruction accuracy and around 8% of the classification accuracy. No other cognitive function was a significant predictor of reconstruction or of classification accuracies. No clear effect of musical expertise was found on reconstruction and classification performances. In conclusion, cognitive factors seem to impact the robustness of the auditory representation and hence the performance of neural based decoding approaches. Taking advantage of this relation could be useful to improve next-generation hearing aids.

## Introduction

The human ability to concentrate attention on one specific sound source in a complex auditory environment is a fundamental skill that allows efficient communication. Indeed, it allows us to follow conversations in crowded places such as classrooms, family dinners or parties. Even if the auditory system seems to be able to perform this task almost effortlessly, concentrating attention on one sound source remains a challenging task that requires a variety of cognitive processes. Such a challenge is magnified for people suffering from profound hearing loss. This is mainly due to the fact that cochlear implants (CI) suffer from a poor spectral resolution (Zeng et al., 2008) and do not allow selective amplification of the source of interest, yielding to a rather poor auditory perception in a noisy environment or in presence of simultaneous auditory sources (Moore., 2003 ; Crandell., 1991 ; Humes et al., 1996). Moreover, it has been hypothesized that the early auditory deprivation may lead to cognitive dysfunctions, also contributing to a poor speech-in-noise comprehension and auditory scene analysis. Therefore, it is particularly important to find a way to either improve auditory scene analysis abilities in CI users or more directly, to improve CI’s amplification systems.

To address this issue, a new approach, called M/EEG-based auditory attention detection (AAD), emerged a few years ago. This approach is based on several works that have demonstrated the possibility to detect which auditory stream, within a set of multiple concurrent streams, an individual was attending to using neural data recorded via electroencephalography (EEG) or magnetoencephalography (MEG) (Ding and Simon, 2012; O’Sullivan et al., 2015; Akram et al., 2014). This relies on the fact that the brain tracks the amplitude envelope of auditory stimuli at the cortical level (Mesgarini et al., 2009 ; Ding and Simon, 2012, Giraud & Poeppel., 2012 ; Nourski et al., 2009; Kubanek et al., 2013). Moreover, in a dichotic context (i.e. different sound in each ear), the cortical tracking of the attended source is enhanced compared to the unattended source (O’Sullivan et al., 2015 ; Pasley et al., 2012; Zion Golumbic et al., 2013). Therefore, these findings are of particular interest to develop neuro-steered hearing aids that are able to detect the source of interest and subsequently amplify it.

However, the literature shows a rather high inter-individual variability in the AAD performance. This variability can be partly explained by physiological factors such as the cranial thickness, the cortical orientation with respect to the skull or the signal to noise ratio of neural data. Nevertheless, it could also be explained by behavioral and cognitive factors including stimulus familiarity, attention, motivation, fatigue and executive functions (Ciccarelli et al., 2019).

Undoubtedly, the ability to selectively listen to a source in a multi-source environment relies upon several cognitive functions. At the outset, focusing attention on one source requires effective segregation of the distinct sources in the auditory scene implying both bottom-up and top-down mechanisms (Bregman, 1994 ; Kondo et al., 2017). The formation of sound streams is particularly dependent on attention because it tends to force the auditory system to group sound elements in line with behavioral or perceptual goals (Shamma et al., 2011, Sussman., 2017, Loui & Wessel., 2007, Snyder et al., 2006). Beside stream segregation per se, it has been shown that attention is important for speech in noise comprehension (Wild et al., 2012). But other, yet related, executive functions seem to be crucial in tasks that require maintaining attentional focus on a source in a cocktail-party scenario.

Firstly, the process of maintaining attention on a task or an auditory object during a certain amount of time is a critical skill for auditory scene analysis and partly depends on sustained attention (Gadea & Espert., 2009 ; Esterman & Rothlein., 2019 ; Tierney et al., 2019). Indeed, sustained attention abilities are linked to better speech in speech comprehension (Tierney et al., 2019) and are also involved in dichotic listening (Helge Johnsen et al., 2002; Hommet et al., 2010; Asbjornsen & Hugdahl., 1995; Hugdahl & Andersson., 1986). However, sustained attention is closely related to other functions and concepts such as selective attention, working memory, inhibitory control, vigilance or motivation (Esterman & Rothlein., 2019). Therefore, it seems to be more a construct than a single cognitive function.

Secondly, working memory (WM) is also important because it is related to speech in noise perception (Besser et al., 2013 ; Sörqvist & Ronnberg., 2012 ; Souza et al., 2015, Escobar et al., 2020), dichotic listening (Conway et al., 2001 ; Colflesh & Conway., 2007 ; James et al., 2014) and in polyphonic music listening (Janata et al., 2002). Additionally, it also seems to play a role in the segregation of streams (Goll et al., 2012) and in the representation of auditory objects (Christison-Lagay et al., 2015) probably because these processes are achieved in the temporal domain. Finally, WM is related to involuntary attentional switching in high cognitive load conditions (Berti & Schröger., 2003).

Finally, inhibitory control seems to be essential to selectively listen to a source in a multi-source environment because inhibition of concurrent streams is necessary. Actually, attentional as well as response inhibition have been shown to be involved in dichotic listening task (Hugdhal et al., 2009 ; Falkenberg et al., 2011), speech in noise (Dryden et al., 2017 ; Knight & Heinrich., 2017 ; Campbell et al., 2020) or auditory scene analysis (Goll et al., 2012). Additionally, in cocktail-party-like scenarios, the different sources can act as distractors and capture attention. Not being able to resist distractions can then lead to involuntary attentional switches resulting in the loss of the source of interest. Moreover, when the cognitive load is high, re-focussing on a target source can be effortful or even impossible, due to the limited amount of attentional resources available (Navon, 1984).

In light of all these elements, WM, sustained attention as well as attentional inhibition could impact performances of AAD algorithms because they support the cognitive processes required when listening to complex auditory scenes.

Yet, another factor could influence the performance of AAD. Actually, in a very recent study, using monodic synthetic musical stimuli, Di Liberto and colleagues demonstrated that average reconstruction performances of their AAD were higher for musical experts than participants with no musical expertise (2020b). This result corroborates previous findings showing a better cortical tracking of auditory stimuli for musicians compared to non musicians (Doelling and Poeppel., 2015 ; Puschmann et al., 2019). Musicians are known to be better in music related tasks such as pitch discrimination (Kishon-Rabin et al. 2001; Micheyl et al. 2006; Magne et al., 2006; Bianchi et al., 2016) or timing perception (Rammsayer & Altenmüller, 2006). They are also better in stream segregation (Zendel & Alain., 2009; François et al. 2014), speech in noise listening (Parbery-Clark et al., 2009a; Coffey et al. 2017) and tend to have better performances in dichotic listening (Barbosa-Luiz et al., 2021). In addition, some works have shown better scores in musicians than in non musicians for cognitive functions involved in cocktail-party like tasks, including cognitive control (Bialystok et al., 2009; Moreno & Farzan, 2015; Moussard et al., 2016 ; Kaganovich et al., 2013), WM (George & Coch., 2011; Jakobson et al., 2008; Zuk et al., 2014, see Talami et al., 2017 for a review) and attention (Medina & Barraza., 2019 ; Strait et al., 2010; Strait & Kraus., 2011; Strait et al., 2015, Clayton et al., 2016).

The first objective of this study was to demonstrate the possibility of reconstructing highly ecological polyphonic musical excerpts from EEG-recorded neural data using a stimulus reconstruction approach and investigating whether neural data allow detecting the focus of attention in a dichotic music listening context. The second and most important goal was to estimate to what extent sustained attention, WM and attentional inhibition can explain the inter-individual variability of AAD in terms of reconstruction and classification performances. Finally, the third objective was to explore whether musicians are better than non musicians in terms of both monodic and dichotic AAD performances.

## Material and Methods

### Participants

A total of forty-five subjects took part in the experiment (28.93 ± 10.94 years, min = 18 years, max = 54 years, 22 females). This number was determined apriori, based on previous studies with the same approach (e.g. O’Sullivan et al., 2015 ; Di Liberto et al., 2021; Cantisani et al., 2019). Over these forty-five participants, one did not complete the online cognitive tests and was thus excluded from the regression analysis, one did not terminate the listening task, and three were discarded from the analysis due to issues related to data acquisition.

All the participants included in the analysis reported no history of hearing disorders, attention impairment or neurological disorder and were not under medication.

Twenty one participants reported having taken formal musical training and two declared being self-taught. Most musicians reported more than 15 years of music making (mean=16.08, sd=5.4) and daily practice. For eight of them, the piano was their main instrument. Four played the guitar and the others played the flute, french horn, harp, saxophone, trombone, trumpet or double-bass. For musicians the mean age was 34.6 ± 13.6 years and 11 of them were females while for non musicians the mean age was 24.7 ± 6.6 years and 9 of them were females.The experiment was approved by the Comité Opérationnel d’Évaluation des Risques Légaux et Éthiques (COERLE) of INRIA and was undertaken in accordance with the Declaration of Helsinki. Each participant provided written informed consent and was financially compensated.

### Main Experimental Task

#### Stimuli

Monodic stimuli consisted of 60 second recordings of four well-known and four less-known classical instrumental piano pieces. The well-known musical excerpts were taken from *Beethoven’s Für Elise, Bach’s Prelude in C Major, Mozart’s Sonata in C Major K545* and *Beethoven’s Pathetique Sonata*. The less-known musical excerpts were taken from *Beethoven’s Lustig und Traurig, Scarlatti Sonata in A Minor K54, Beethoven’s Sonata in F Presto* and *Bach’s Prelude no°8*.

Dichotic stimuli consisted of pairs of musical excerpts simultaneously presented to the left and the right ear. Pairs were composed of rhythmically similar musical excerpts as assessed via Power Spectrum Density measures (see Figure 1). This allowed to ensure that the attention would not be systematically more captured by a greater presence of notes in one excerpt of the pair. The pairs were *Beethoven’s Fur Elise/Bach’s Prelude in C Major, Mozart’s Sonata in C Major K545/Beethoven’s Pathetique Sonata, Beethoven’s Lustig und Traurig/Scarlatti’s Sonata in A Minor K54* and *Beethoven’s Sonata in F Presto /Bach’s Prelude no°8*. All musical excerpts were RMS normalized at the same amplitude. Stimuli were presented via Sennheiser HD-25 supra-aural headphones at a comfortable level.

**Figure 1:**
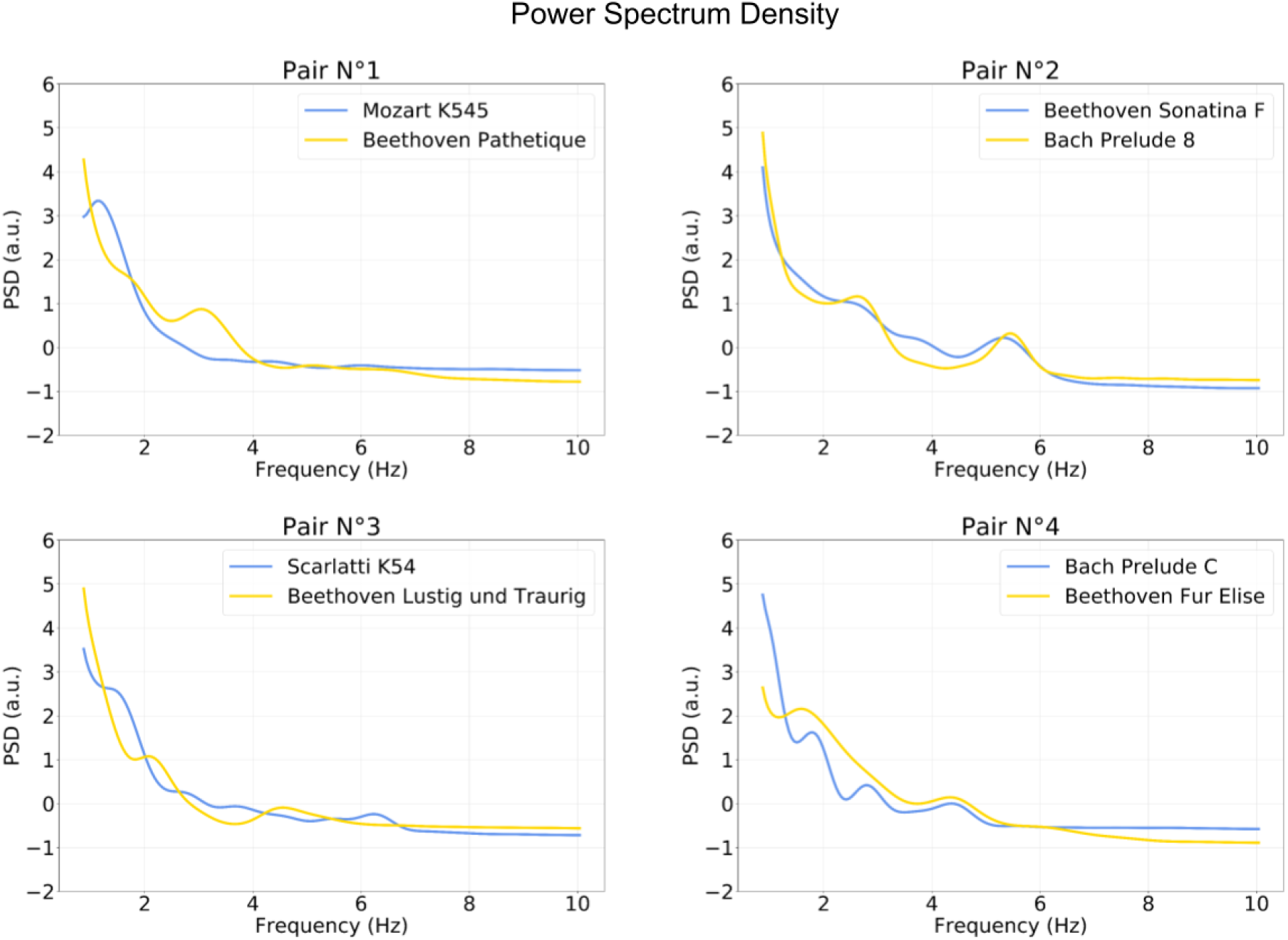
Power Spectrum Density plots of the envelope of each musical excerpt. The excerpts were presented either individually in the monodic condition or by pair in the dichotic condition. Pairs were chosen in order to minimize differences in the rhythmic structure.

#### Procedure

Participants were comfortably seated in front of a 28’ computer screen at a distance of approximately 80 cm. The task was divided in two parts : Monodic and Dichotic.

In the monodic part, participants were instructed to concentrate their attention on the musical excerpt as if they were trying to memorize it. Then, the musical excerpt was played for 60 seconds. During the stimulus presentation, a fixation cross was displayed at the screen center. Each of the musical excerpts was presented twice for a total of sixteen 60 second trials. The order of presentation avoided direct repetition of a given excerpt.

In the dichotic part, four pairs of musical excerpts were presented dichotically (±90° from the azimuth). Participants were visually instructed which side (left or right) they had to attend. Each pair was presented four times (attend left, attend right for each of the two excerpts in the pair) for a total of sixteen 60 second trials. The whole experiment was monitored using a custom version of OpenSesame developed by Oticon Medical (Omexp : Patou et al., 2019).

### Cognitive Tasks

In this study we were interested in three cognitive functions that are closely related to auditory scene analysis : WM, Attentional Inhibition and Sustained Attention. Attentional inhibition and sustained attention were measured online and from home for every participant using OpenSesame (Mathôt et al., 2012) and JATOS online test platform (Lange et al., 2015). This remote testing prevented an excessively long experimental session.

#### Reverse Digit Span Task

To measure WM, we used the Reverse Digit Span Task (RDST). The RDST is a classical WM task in which participants are presented with an auditory sequence of digits (between 1 and 9) that they have to immediately report orally in the reverse order. The length of the sequence increases until the participant makes two errors in a row. The test starts with a sequence of three digits and can reach a maximum of nine digits. The WM score corresponds to the size of the longest correctly reported sequence. The RDST was performed just before the listening task.

#### Auditory Stroop

We measured attentional inhibition with an online auditory Stroop task adapted from Donohue et al (2012). The auditory stimuli consisted of French words “aigu” (“high”) and “grave” (“low”) spoken by a native French speaker, digitally sampled at 44100 Hz and resynthesized (Praat : Boersma, 2001) to obtain two tokens of each word with an average fundamental frequency of ∼80 and 260Hz. All tokens lasted 300ms. Congruent trials consisted of the word “aigu” pronounced in a high pitch and the word “grave” pronounced in a low pitch while incongruent trials consisted of the word “aigu” pronounced in a low pitch and the word “grave” pronounced in a high pitch. Participants were instructed to decide as quickly and accurately as possible what was the pitch of the spoken word, regardless of its meaning, by pressing on a keyboard. The response side to low or high pitch was counterbalanced across participants. The SOA was set to 1350 ms plus the time needed to respond.

A total of 260 congruent and 260 incongruent trials were presented in a pseudo-random order in10 blocks of 52 trials (20 minutes).

For each participant we computed the accuracy (% of correct responses), the reaction times, the efficiency score for congruent and incongruent trials (accuracy/RTs) and an Inhibition score (Congruent efficiency -Incongruent efficiency). Thus, the higher the inhibition score, the better the attentional inhibition. The computation of these metrics allow one to take into account the Speed Accuracy Trade Off (SATO) (Gartner & Strobe, 2020).

#### Auditory Sustained Attention To Response Task (SART)

We measured sustained attention with an online auditory SART adapted from Seli et al (2012). The auditory stimuli consisted of digits (from 1 to 9) spoken by a native French speaker digitally sampled at 44100 Hz. Their duration was ∼ 400 ms and they were randomly presented. Each digit was followed by a pink noise mask for 900ms. Participants were instructed to press the spacebar as quickly and accurately as possible each time they heard a digit except when they heard the digit 3. In this case, they had to withhold their answer. A total of 225 trials (200 Go and 25 NoGo) were undertaken by the participants, lasting around 7 minutes.

Accuracy for Go and NoGo trials were computed as well as RTs for Go trials. A sustained Attention score was computed as follows : 1000 * (Accuracy_NoGo_Trials / Average_RT_Go_Trials). The higher the sustained attention score, the better the sustained attention. As for attentional inhibition the computation of this complex metric allows one to take into account the SATO in SART performances (Seli et al., 2013 ; Seli et al., 2016 ; Hallion et al., 2020).

### EEG Data

Electroencephalography data were recorded using an ANT Refa8 amplifier and a 21-electrode cap (arranged in accordance with the International 10-20 system) with a sampling rate of 256 Hz and an average reference. They were preprocessed using the *MNE-python* package (Gramfort et al., 2013) according to the procedure described in Crosse et al., 2021. First, raw EEG data were cut into 60s epochs, each epoch representing a trial, and the first 500ms of each epoch was discarded to avoid modeling the response to the stimulus onset. Next, EEG data were digitally filtered between 1-40Hz and bad channels were interpolated via spherical spline interpolation when necessary. Then, ICA was performed to remove eye-blinks and saccades artifacts. Afterward, EEG data were digitally filtered between 1-9Hz (Greenlaw et al., 2020 ; Biesman et al., 2015) using a 4th order Butterworth zero-phase-shift filter and downsampled to 64Hz. Finally, 60s epochs were cut into 30s epochs, resulting in 32 epochs for monodic and dichotic conditions, with the aim of having enough data to train and test the stimulus-reconstruction model.

### Audio Feature Extraction

Amplitude envelopes of the audio monodic stimuli were obtained using the function *human_cochleagram* from the Python *Pycochleogram* package (https://github.com/mcdermottLab/pycochleagram). This function allows 1) computing an Equivalent Rectangular Bandwidth (ERB) filter bank and 2) using this filter bank to decompose the signal into subband envelopes. Afterwards subband envelopes were averaged to obtain a unique envelope. Each envelope was then digitally filtered between 1-9Hz with a 4th order Butterworth zero-phase-shift filter, downsampled to 64Hz and cut into 30s long epochs in order to match EEG preprocessing.

### Stimulus Reconstruction

One objective of this study was to investigate if it was possible to reconstruct and classify monodic and dichotic natural polyphonic music from neural data. To do so, we used the same stimulus reconstruction approach as O’Sullivan et al. (2015) (this approach is also described in several other articles, see Crosse et al., 2016 and Crosse et al., 2021 for a comprehensive description).

#### General procedure

The stimulus reconstruction approach allows one to reconstruct an estimate of the envelope of the auditory stimulus *s* using electrophysiological neural data *d* via a linear reconstruction model *g* (O’Sullivan et al., 2015). The reconstruction model *g(τ, n)* is a temporal response function (see Lalor et al., 2009 for details) that maps neural data *d(t, n)* to stimulus *s(t)* as follow :

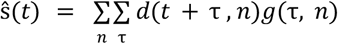

Where *ŝ* denotes the reconstructed stimulus, *d(t,n)* represent the response of electrode n at time t=1 … T and *τ* are some time lags that represent a window in which the brain’s response to the stimulus is supposedly optimal. We defined *τ* to go from 0.200ms pre-stimulus to 0.350ms post-stimulus based on previous works (O’Sullivan et al., 2015, Di Liberto et al., 2020b, 2021). At a sampling rate of 64 Hz, this corresponds to 36 sample shifts (incl. the zero shift).

The model g is estimated by minimizing the mean-squared error between the original stimulus *s(t)* and the reconstructed one *ŝ (t)*:

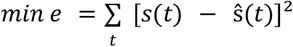

A robust minimizer of the mean-squared error is obtained using the following matrix operations:

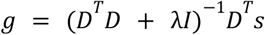

Where *D* is the lagged time series of the response matrix *d* and *λ* is a ridge parameter term introduced to avoid overfitting. The model parameters as well as the ridge parameter are generally estimated using leave-one-out cross validation (LOO) procedure. Once the model parameters have been tuned, it can be tested on new data (generally the leaved out fold) and the end of this process is the reconstructed stimulus *ŝ*. Finally, a reconstruction score is computed, to estimate the reconstruction accuracy of the model, as the Pearson correlation between the reconstructed stimulus ŝ and the original stimulus *s*. The higher the reconstruction score, the better the reconstruction accuracy.

##### Training and testing the model

In this study, we wanted to see if we can reconstruct a trial with a model trained on every other monodic/dichotic trial. To do so, we used a custom Python script where a first LOO was performed to separate training and test phase and because we introduced a ridge parameter (*λ*) we used a second LOO nested in the training phase to estimate the optimal value of the ridge parameter from a set of 20 logarithmically spaced values from ^−10^to ^10^.

At the end of the test phase, a reconstruction s_i_ is obtained for each musical excerpt s_i_, based on a model trained on all the other musical excerpts. This procedure allowed us to make sure that the musical excerpt we wanted to reconstruct was only used to validate the model.

Because we had 32 monodic and 32 dichotic trials, we trained separate models for each trial type. Therefore, this resulted in 32 models for monodic trials for each participant. For the dichotic trials, unlike O’Sullivan et al. (2015), we chose to train the models to reconstruct the attended musical excerpts only because 1) we were not interested in estimating which musical excerpt the participant was not attending to and 2) their results indicate that performances of unattended models are worse than performances of attended models. Thus, this resulted in 32 dichotic models per participant.

#### Performances Metrics

##### Reconstruction Accuracy

The reconstruction accuracy corresponds to the Pearson correlation between the reconstructed envelopes and the original musical envelopes. This led to 32 r values per participant. An average reconstruction accuracy score (*r*_*mono*_) was also computed per participant.

Similarly, for dichotic trials, the reconstruction was computed for both attended (*r*_*Attended*_) and unattended (*r*_*Unattended*_) musical excerpts as the Pearson correlation between the reconstructed envelopes and the original attended and unattended musical envelopes.

Because we were interested in evaluating the increase in performance for attended vs the unattended stimuli, we defined a Dichotic Performance Index (DPI) as follows :

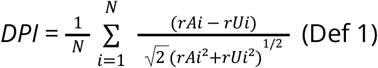

Where *rA* and *rU* are the *r*_*Attended*_ and *r*_*Unattended*_ scores of each trial. The advantage of this measure is to provide for each participant a single measure summarizing the distance between attended and unattended r values while being robust to inter-trial variability.

##### Classification Accuracy

Aside from reconstruction accuracy, we computed a classification accuracy score. For monodic trials, the classification accuracy score was the percentage of monodic trials that were correctly identified. We considered a trial to be correctly identified when *r*_*mono*_ was higher than all the correlations between the reconstructed musical excerpts envelope and the other musical excerpts envelope. Similarly, for dichotic trials, we considered a trial to be correctly identified when *r*_*Attended*_ was higher than *r*_*Unattended*_. This score reflects the percentage of trials for which we were able to correctly classify the attended/unattended music, in dichotic trials.

## Statistical Analysis

All statistical analyses were performed using the R software (R Core Team, 2022) with the lme4 (Bates et al., 2015), stats (R Core Team, 2022), car (Fox, 2019) and lmerTest (Kuznetsova et al., 2017) packages.

### Normality testing, Outliers Rejection and Effect sizes

When appropriate, we assessed normality of the data distributions with a Shapiro-Wilk test. We tested for potential outliers using either a Grubbs test or a Tukey Fence for Outliers. We assessed heteroscedasticity using either an F test or a Fligner test. Finally, to measure the effect sizes we computed either a Cohen’s d or a Rank-Biserial Correlation.

### Cognitive Functions

#### Online test reliability

To test the reliability of online cognitive tasks, we compared Go/Congruent and NoGo/Incongruent distributions using a Wilcoxon rank sum test as differences between the two classes were strongly predicted by previous findings (REFs).

#### Effect of musical expertise

To see whether musical expertise has an impact on cognitive functions, we compared the cognitive function test’s performance of participants with low and high musical expertise using either a Student’s t test or a Wilcoxon rank sum test, as appropriate.

#### Performance Metrics

##### Individual and group chance levels for classification accuracy

Individual, as well as group classification accuracy chance levels were estimated with an inverse cumulative probability distribution function for binomial distribution (O’Sullivan et al., 2015) and computed using the R *qbinom* function.

##### Effect of musical expertise

To test whether musical expertise has an impact on reconstruction and classification accuracies we, again, compared the distributions of participants with low and high musical expertise using Student t test as they were all normal.

Additionally, to see if the distributions of participants with low and high musical expertise were different from the null distribution for reconstruction and classification accuracies, we used a One sample Student t test, with μ = 0 and with μ = 50, respectively.

#### Cognitive Functions and Performances metrics

In order to see if the chosen cognitive functions were good predictors of the reconstruction and classification accuracies, we used linear models computed on averaged data. A backward stepwise regression, where all the predictors (i.e. sustained attention, WM and attentional inhibition) were entered simultaneously, was used to select the best model for each dependent variable (i.e. reconstruction and classification accuracies) in each condition (monodic and dichotic). The best model was selected as the one with the lowest Akaike Information Criterion (AIC). After the best model was selected, models’ assumptions of normality, homoscedasticity and independence of the residuals were confirmed using Jarque-Bera, Breush-Pagan and Durbin-Watson tests respectively. Finally, effect sizes were computed as the Cohen’s f2.

## Results

The objectives of this study were: 1) to test if it was possible to reconstruct and classify monodic and dichotic natural polyphonic music from neural data, 2) to investigate the relation between specific executive functions and the reconstruction and classification performances and 3) to explore the impact of musical expertise on these above-mentioned performances.

First, some results about cognitive functions are briefly exposed. Then, results about reconstruction and classification performances for both monodic and dichotic stimuli are presented. Next, results about the relation between executive functions and reconstruction and classification performances are exposed. Finally, the impact of musical expertise over the performances are reported.

### Cognitive Functions

Because we used an online version of the SART to measure sustained attention, we first compared the accuracy for Go and NoGo trials in order to see whether we could reproduce the findings reported in the literature. After outlier correction (see Methods), the test indicates a very large difference between the accuracy of Go (M=99.3% sd=±1.65%) and NoGo trials (M=77% ±14.8%) (*W = 1346.00, p < .001; r = 0.97*). This confirms the reliability of the online version of the SART.

We proceeded similarly with the data of the online auditory Stroop test, by comparing the global efficiency of congruent and incongruent trials. The test indicates a large positive difference between global efficiency of congruent (M=166.7 ±28.5) and incongruent trials (M=151.9 ±24.4) (*W = 1040.00, p = 0.021; r = 0.30*). This confirms the reliability of the online version of the auditory Stroop task.

Then, we explored whether musical expertise has an impact on the selected cognitive function, by comparing participants with low (N=21) and high (N=19) musical expertise. For WM, the effect of musical expertise was non-significant (M-low=5.04 sd=±1.2, M-high=5.47 sd=±1.2, *w=238, p = 0.287, r = 0.19*).

For sustained attention scores, the effect of musical expertise was also non-significant (M-low=149.83 ±38.2, M-high=165.39 ±32.7, *t(38)=1.37, p = 0.17, Cohen’s d = 0.44*).

Finally, for attentional inhibition scores, the effect of musical expertise was again non-significant (M-low=13.6 ±11.8, M-high=16.3 ±11.4, *t(38)=0.73, p = 0.47, Cohen’s d = 0.23*). Thus, overall, musical expertise did not have a significant effect on the cognitive variables we measured, although one can observe that values tended to be greater for participants with high compared to low musical expertise (see Figure 2).

**Figure 2:**
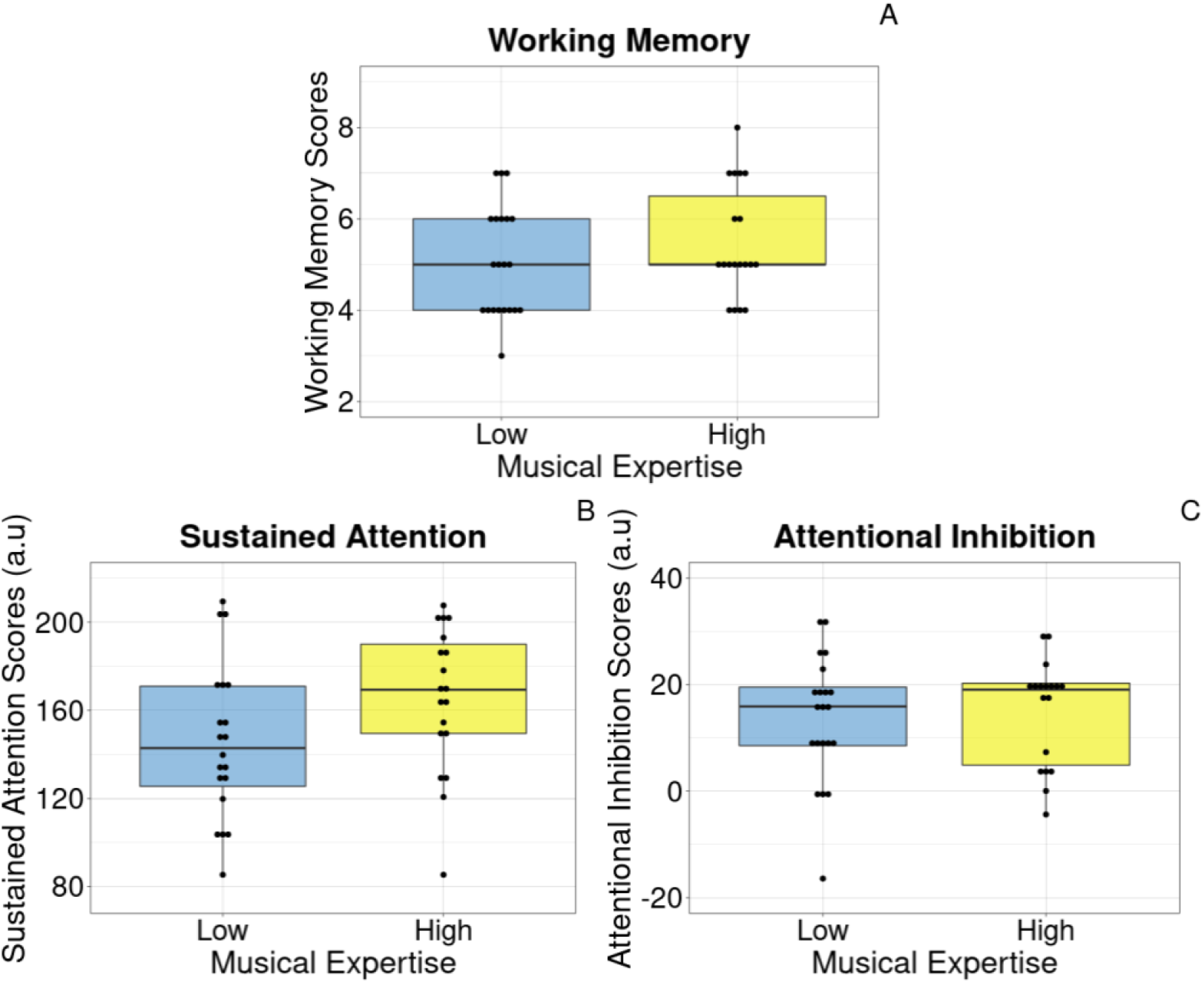
Effect of musical expertise on cognitive functions. (A) WM scores for participants with low musical expertise (left) and high musical expertise (right). (B) Sustained Attention scores for participants with low musical expertise (left) and high musical expertise (right). Greater values indicates better sustained attention ability. (C) Attentional inhibition scores for participants with low musical expertise (left) and high musical expertise (right). Greater values indicate better attentional inhibition. The central line of the box plots represents the median of the distribution, the lower and upper hinges correspond to the first and third quartiles and the whiskers represent the largest and lowest values no further than 1.5 times the interquartile range from the hinges. Individual points represent individual data.

### AAD Performance

#### Reconstruction Accuracy

One of the objectives of this study was to test if one can reconstruct monodic and dichotic natural polyphonic music from neural data, with a simple backward linear model.

On average, the reconstruction performance of the monodic musical excerpts was 0.063 (± 0.08). This result is significantly different from the null distribution (*t(40) = 14.32, p < 0.001* ; *Cohen’s d = 2.24*). Moreover, the average correlations between the reconstructions of the target excerpts (i.e. the listened excerpts) and their corresponding original envelopes were higher than all the correlations between the reconstructions of the target excerpts and the other original envelopes, as indicated by the higher values on the reconstruction matrix diagonal (figure 3 A).

**Figure 3:**
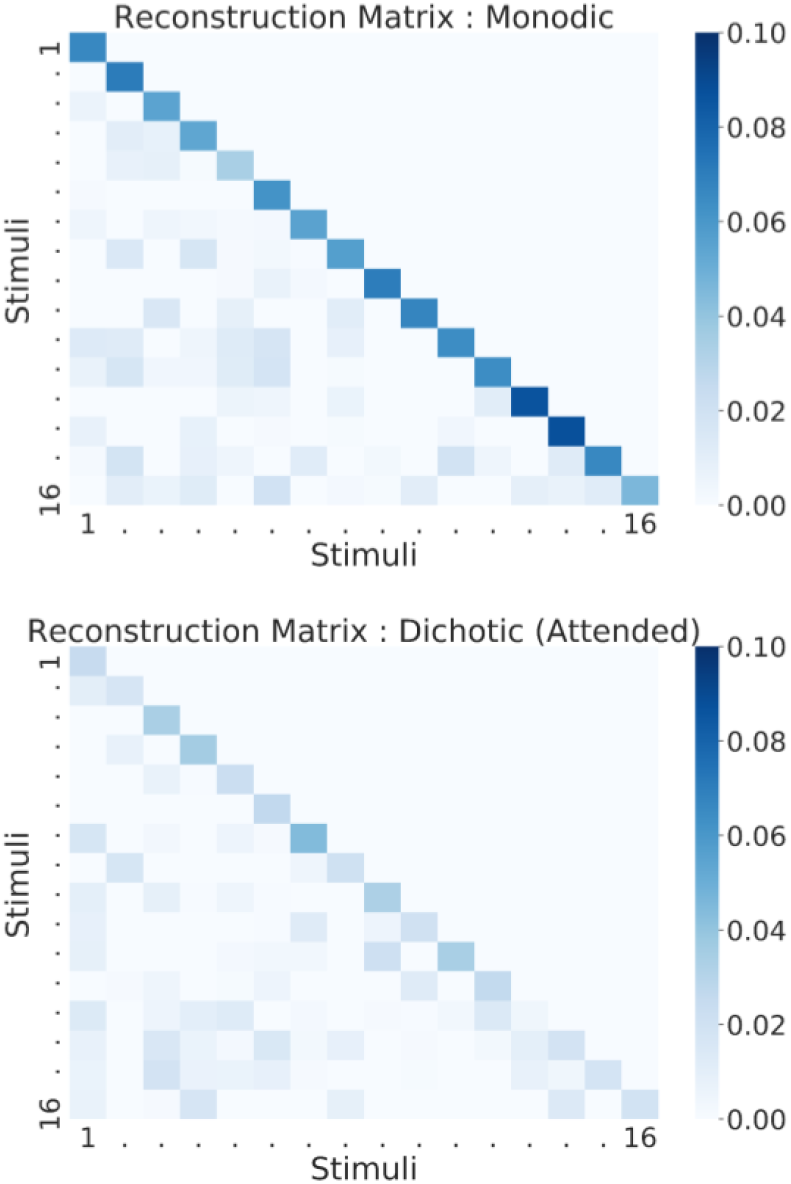
Heatmaps of reconstruction scores for monodic (A) and dichotic (B) stimuli. Values on the diagonal indicate the reconstruction of the musical excerpt with the corresponding original envelope (e.g. correlation between the reconstruction of Beethoven Fur Elise and the original envelope of Beethoven Fur Elise). In (B) values on the diagonal indicate the reconstruction of the attended musical excerpt with the corresponding original envelope. Darker colors indicate higher reconstruction values. Reconstruction scores were averaged across repetitions and participants. Overall, for both monodic and dichotic attended stimuli, reconstruction values are higher when computed with the original envelope rather than with the envelope of another stimulus.

For the dichotic stimuli, the average reconstruction performance was 0.028 (± 0.018) for the attended musical excerpts, 0.013 (± 0.013) for the unattended excerpts and 0.1 (± 0.16) for the DPI (Def 1). These reconstruction performances were clearly different from the null distribution : rAttended (*w=856, p < 0.001* ; *r = 0.99*), rUnattended (*w=749, p < 0.001* ; *r = 0.83*). Importantly, attended streams were better reconstructed than unattended ones (*w=1191, p < 0.001* ; *r = 0.45*). Moreover, for attended streams the average correlations between the reconstructions of the target excerpts (i.e. the attended excerpts) and their corresponding original envelopes were higher than all the correlations between the reconstructions of the target excerpts and the other original envelopes, as indicated by the higher values on the reconstruction matrix diagonal (see figure 3 B).

Figure 4 shows a great inter-individual variability in the reconstructions of both monodic and dichotic musical excerpts.

**Figure 4:**
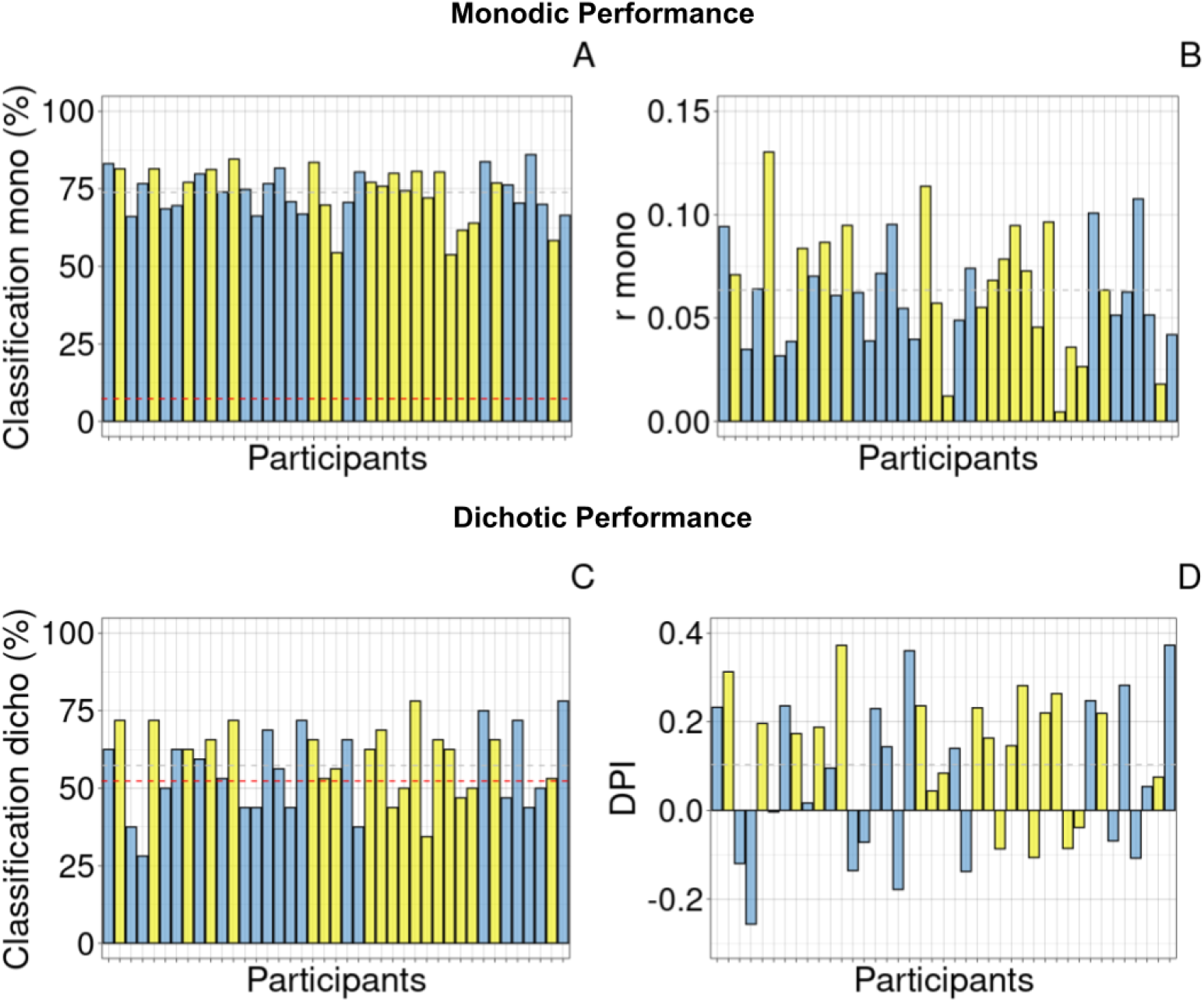
Illustration of monodic and dichotic reconstruction and classification accuracy. (A) Classification Performance for monodic stimuli (average = 73.8%). (B) Reconstruction performances of monodic stimuli (average = 0.063). (C) Classification Performance for dichotic stimuli (average = 57.31%, group chance level = 52.3%). (D) Dichotic Performance Index (DPI) of dichotic stimuli (average = 0.102), representing the distance between attended and unattended r values. Gray dashed lines indicate average reconstruction and classification. Red dashed lines indicate the group chance level. Blue-filled bars represent participants with low musical expertise while yellow-filled bars represent participants with high musical expertise.

We were also interested to see whether the monodic reconstruction performance could explain the dichotic one (Figure 5). Therefore, we computed a Pearson correlation that indicates a non-significant relation between the two performances (r = 0.2, p-value = 0.197).

**Figure 5:**
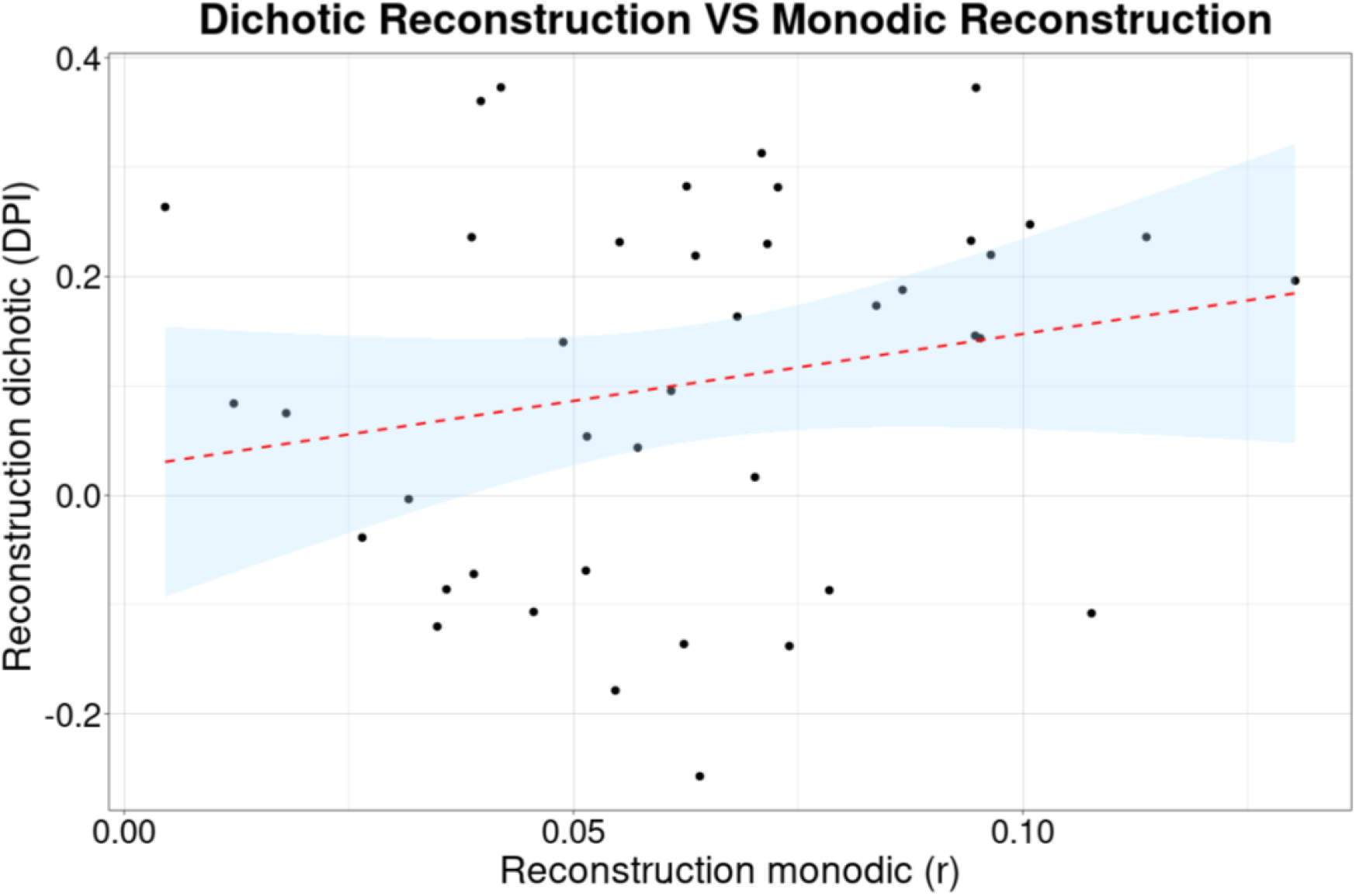
Scatterplot illustrating the lack of significant relation between dichotic and monodic reconstruction. Adjusted-R^2^ = 0.017, p-value = 0.2. Red dashed line indicates the regression line. Blue area indicates confidence interval at 95%.

#### Classification Accuracy

In addition to the reconstruction accuracy, we also investigated to what extent we could correctly classify the stimuli (Figure 3).

On average, classification accuracy for monodic trials reached 73.8% (± 8.22) and all the participants had a classification accuracy significantly greater than chance level (=12.5%, estimated using an inverse binomial test ; see method). At the group level the average classification accuracy was clearly higher than the group chance level (=7.3%, estimated using an inverse binomial test).

As expected, the classification accuracy of dichotic trials was lower and reached 57.31% (± 12.8). At the individual level, only 10 participants out of 41 had a classification accuracy significantly greater than chance level (=65.65%, estimated using an inverse binomial test). However, at the group level, the average classification accuracy was higher than the group chance level (=52.3%, estimated using an inverse binomial test).

### Effect of Musical Expertise on Reconstruction and Classification Accuracies

We were also interested in knowing whether the degree of musical expertise could have an impact on both reconstruction and classification accuracy of monodic and dichotic stimuli (Figure 6).

**Figure 6:**
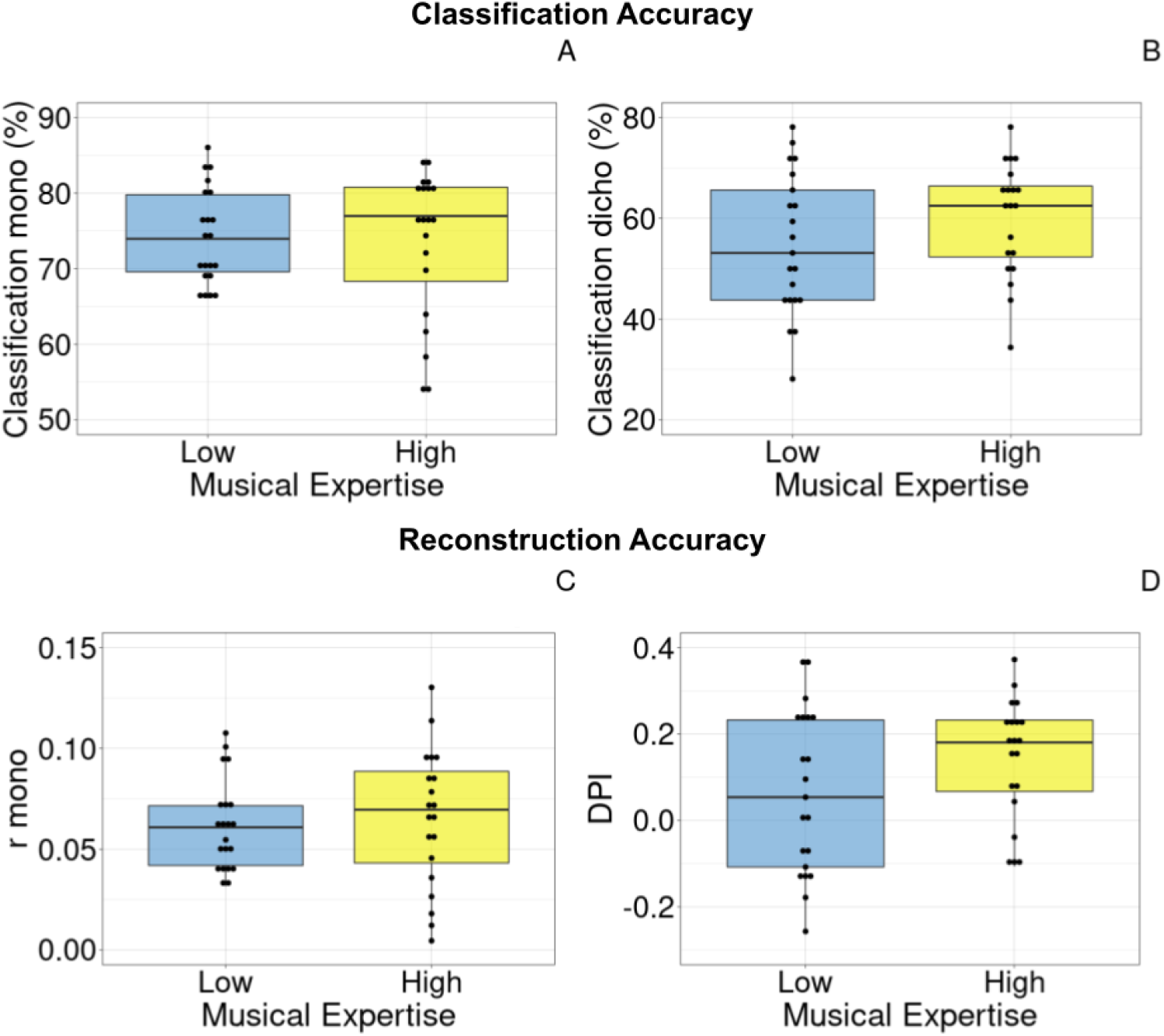
Effect of musical expertise on monodic and dichotic reconstruction and classification. (A) Classification of monodic stimuli for low (left) and high musical expertise (right). (B) Classification of dichotic stimuli for low (left) and high musical expertise (right). (C) Reconstruction of monodic stimuli for low (left) and high musical expertise (right). (D) DPI of dichotic stimuli for low (left) and high musical expertise (right). The central line of the box plots represents the median of the distribution, the lower and upper hinges correspond to the first and third quartiles and the whiskers represent the largest and lowest values no further than 1.5 times the interquartile range from the hinges. Individual points represent individual data.

We found a clear difference between the null distribution (μ = 50) and both the DPI and dichotic classification accuracy of high musical expertise participants : DPI (t(19)=4.64, p < 0.001 ; Cohen’s d = 1.04), Dichotic classification accuracy (t(19)=4, p < 0.001 ; Cohen’s d = .89). However, we found no difference between the null distribution and the dichotic performances of low musical expertise participants : DPI (t(20)=1.54, p = 0.14, cohen’s d = 0.34), Dichotic classification accuracy (t(20)=1.56, p = 0.13 ; cohen’s d = 0.34).

However, the direct effect of musical expertise was significant neither for monodic (M-low=0.061 ±0.022, M-high=0.065 ±0.034, *t(39)=0.42, p = 0.67* ; *Cohen’s d = 0.13*) nor for dichotic reconstruction (M-low=0.063 ±0.18, M-high=0.144 ±0.13, *t(39)=1.56, p > 0.12* ; *Cohen’s d = 0.49*) and not either for both monodic (M-low=74.22 ±6.34, M-high=73.42 ±9.8, *t(39)=0.31, p = 0.75* ; *Cohen’s d = 0.1*) nor for dichotic classification (M-low=54.8 ±13.9, M-high=60 ±11.2, *t(39)=1.32, p = 0.19* ; *Cohen’s d = 0.41*).

### Effect of Cognitive Functions on Reconstruction and Classification Accuracies

The main goal of the present study was to test whether selected cognitive functions could explain a part of the inter-individual variability of the reconstruction and classification performances during dichotic listening as well as during the listening of monodic stimuli (Figure 7).

**Figure 7:**
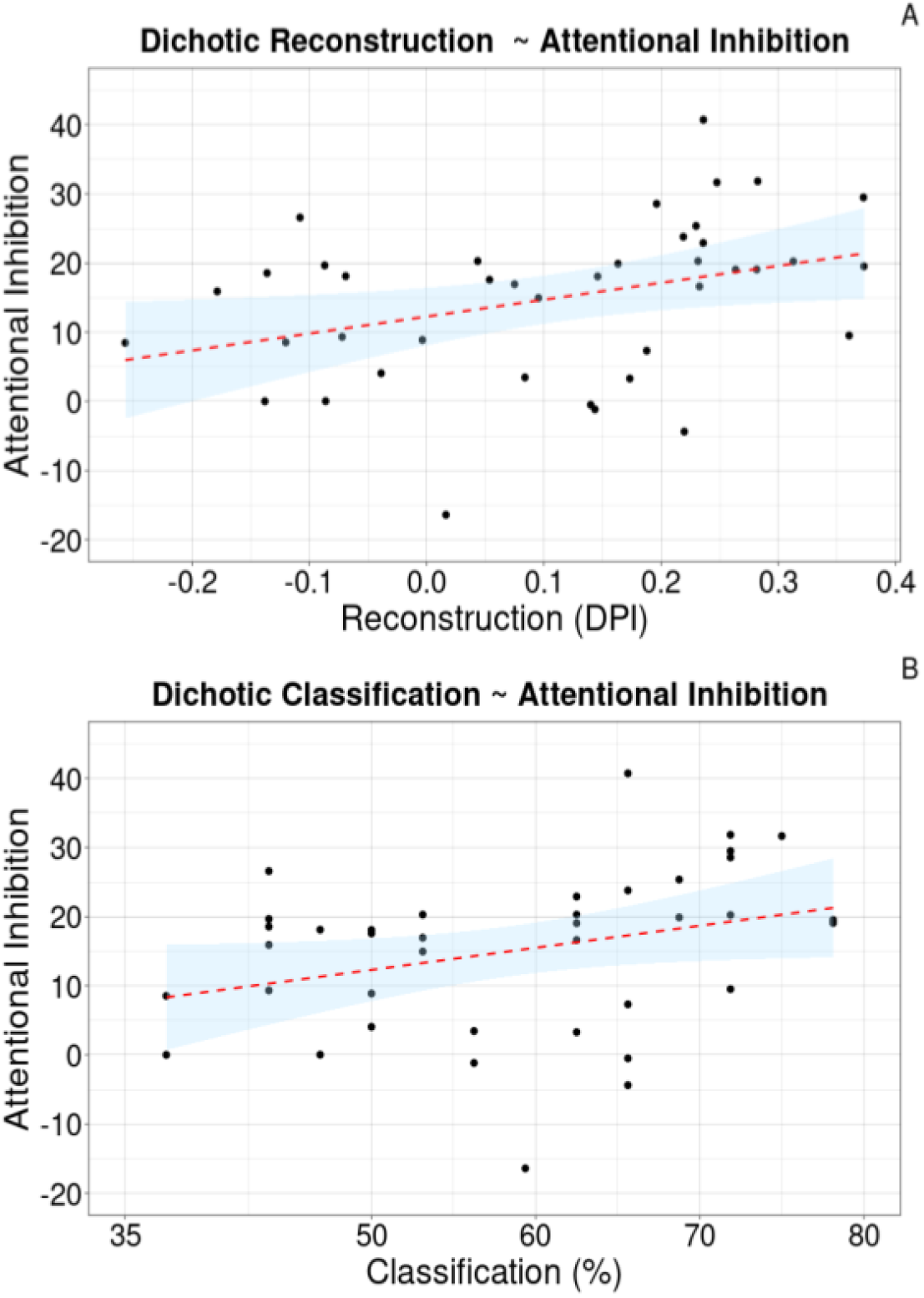
Illustration of the significant relation between attentional inhibition and dichotic listening metrics. (A) Linear regression of the DPI by attentional inhibition. Adjusted-R^2^ = 0.1, p-value = 0.023. (B) Linear regression of the dichotic classification accuracy by attentional inhibition. Adjusted-R^2^ = 0.08, p-value = 0.039. Red dashed line indicates the regression line. Blue area indicates the confidence interval at 95%.

#### Attentional Inhibition

For monodic stimuli we found no effect of attentional inhibition on reconstruction accuracy (*t(38)=1.88, p > 0.068*) and the prediction model was not significant (*adjusted-R*^*2*^ *= .06, f(1,38)=3.52, p = 0.068, Cohen’s f2 = 0.09*). There was also no effect of attentional inhibition on classification accuracy *(t(38)=1.42, p = 0.16*). Once again the prediction model was not significant (*adjusted-R*^*2*^ *= .03, f(1,38)=2.01, p = 0.16, Cohen’s f2 = 0.05*).

For dichotic stimuli, we found a medium positive effect of attentional inhibition on DPI (t(38)=2.36, p = 0.024) and attentional inhibition predicted around 10% of the DPI (adjusted-R=.1046, f(1, 38)=5.55, p = 0.024, Cohen’s f2 = 0.15). This regression model was significantly different from the null model (F(2,39) = 5.55, p = 0.023). We also found a small but significant positive relation between attentional inhibition and classification accuracy (*t(38)=2.13, p = 0.039*). Attentional inhibition predicted around 8% of the classification accuracy (*adjusted-R*^*2*^ *= .0835, f(1,38)=4.56, p = 0.039, Cohen’s f2 = 0.12*). This regression model was significantly different from the null model (F(2,39) = 4.55, p = 0.039).

Additionally, no significant interaction was found between attentional inhibition and any of the other cognitive variables.

#### Other Cognitive Functions

For WM and sustained attention we found no effect on monodic nor dichotic reconstruction (*t(38) < 2, p > 0.05*) nor classification accuracy either (*t(38) < 2, p > 0.05*).

Using a backward stepwise regression (see Material & Methods), we compared a full model with all the cognitive functions as predictors, to a simple model with attentional inhibition only for both dichotic reconstruction and classification accuracy. The AIC criteria of the simple model were lower (DPI = -145.1 and classification = 200) than the one of the full model (DPI = -142.2 and classification = 202.7) suggesting that the full model was not significantly better than the simple one.

## Discussion

In the present study we used a reconstruction stimulus approach to explore whether it was possible to reconstruct and classify both monodic and dichotic natural polyphonic musical excerpts on the basis of EEG recordings. We also measured sustained attention, attentional inhibition and working memory via cognitive tests either online (SART and Stroop tests) or with a more traditional approach (Reverse Digit Span Task). Then, we employed linear regression models to see whether cognitive functions and musical expertise explain reconstruction and classification performance of neural data. Additionally, we explored whether musical expertise affects executive functions as well as neural tracking of musical stimuli by comparing non-musical and musical experts.

First, our results show that it is possible to reconstruct both monodic and dichotic natural polyphonic excerpts based on neural data using the reconstruction stimulus approach (see Figure 3). In particular, for the dichotic stimuli, attended stimuli were better decoded than unattended stimuli. Moreover, it was also possible to classify not only the monodic stimuli with good precision (73.8%), but also the dichotic stimuli although to a lesser extent (57.3%).

Second, unlike what we expected, we found no difference in terms of cognitive functions between participants with high and low musical expertise. Moreover, we found no effect of musical expertise on reconstruction and classification performances for monodic nor dichotic stimuli.

Finally and most importantly, while for monodic stimuli we found no effect of the selected cognitive functions over reconstruction and classification accuracies, for dichotic stimuli attentional inhibition did explain around 10% of the reconstruction accuracy and around 8% of the classification accuracy. No other cognitive function was a significant predictor neither of dichotic reconstruction nor of classification accuracies.

We will first discuss the feasibility using ecological musical stimuli within the auditory attention decoding approach. Then, we will discuss the relation between cognition and performances of stimulus reconstruction. Finally, we will consider the effect of musical expertise over the chosen algorithm performances.

In this work, we explored whether it was possible to both reconstruct and classify monodic and dichotic ecological musical excerpts based on neural data.

In the monodic condition, reconstruction performances were not very high (0.064 on average) but significantly greater than a null distribution. The reconstruction matrix and the resulting diagonal (Figure 2), indicate that the average correlation between the reconstructed musical excerpts and their corresponding original envelopes were clearly higher than the average correlations between the reconstructed musical excerpts and all the other envelopes. This result indicates that, at the group level, no matter their magnitudes, the reconstructions are more similar to the original envelopes than to the other ones. In addition, reconstruction values using polyphonic music stimuli were close to the results obtained in other similar studies using monodic music stimuli (e.g. Di Liberto et al., 2020a), although somewhat smaller than when using multiway canonical correlation analysis (Di Liberto et al. 2020b, 2021). In the dichotic condition, as expected, reconstruction performances were lower (0.028 on average for attended stimuli, 0.013 for unattended ones and 0.1 on average for the DPI). However, reconstruction was significantly greater than chance for both attended and unattended stimuli. Moreover, the greater reconstruction performances for attended compared to unattended stimuli shows that participants were able to do the task, namely concentrate their attention on the target stream and inhibit the concurrent one, despite the reported difficulty to do so. Overall, these results confirm previous findings showing that the brain tracks the amplitude envelope of target and concurrent streams (e.g. Ding and Simon., 2012 and O’Sullivan et al., 2015). Importantly, the replication of these previous findings is a crucial test to confirm that the design and analysis pipeline were appropriate to reconstruct both monodic and dichotic realistic polyphonic musical stimuli based on neural data.

Classification performance can be interpreted in a similar line. In the monodic condition, classification of the attended musical excerpt was good (around 74%) and consistent with previous findings (Schaefer et al., 2011). In addition, all participants showed a classification accuracy clearly higher than chance level (7.3% for a thirty-class problem). For the dichotic condition, the accuracy to classify the target stream was lower (57%), but still higher than chance at the group level (= 52%). This is consistent with the work of Cantisani and colleagues (2019) who found a classification accuracy of around 58% for duet stimuli. However, while their duet stimuli were composed of two different monophonic instruments, we used only pairs of polyphonic piano excerpts. This leads to a more complicated auditory scene in our experiment because 1) in a single excerpt, the piano plays several notes at a time, 2) the timbre of two different pianos is rather similar. Overall, the results discussed here indicate that, using a reconstruction stimulus strategy, it was possible not only to detect which musical excerpt between many was presented in a monodic condition but also to detect which musical excerpt was attended in a rather challenging dichotic listening context.

However, it is important to consider that interindividual variances of the reconstruction and classification performances in both conditions were relatively high, in particular in the dichotic condition. Indeed, the DPI ranged from ∼-0.28 to ∼0.39 and the classification from almost 30% to around 80% in dichotic listening. One possible explanation of such variability could be the different quality of EEG data across participants. Nevertheless, because classification and DPI are not sensitive to the magnitude of the reconstruction values, this variability should probably not be attributed to the data quality. Indeed, classification accuracy is computed as the percentage of trials for which r_Attended_ is higher than r_Unattended._ Moreover, the absolute values of *r*_*Attended*_ and *r*_*Unattended*_ do not affect the computation of the DPI (see Material & Methods section). Therefore, these metrics should not be biased by inter-trial and inter-participant variability caused by data quality fluctuations. This hypothesis is also corroborated by the non-significant correlations between monodic reconstruction and DPI (r=0.2, p-value=0.197, see Figure 5) and between the reconstruction of monodic and both Attended (r=0.22, p-value=0.16) and Unattended dichotic stimuli (r=0.06, p-value=0.66). Indeed, if one considers the monodic condition to be the baseline, then the absence of correlation between the baseline and the experimental condition (i.e. dichotic listening) suggests that factors such as skull thickness, brain orientation or Signal to Noise Ratio are not likely to explain the dichotic performances. Importantly, it also provides hints on the fact that dichotic AAD performance seems to be dependent on other factors than just the general ability to listen to a musical piece.

One possible factor that we assessed in the present work is musical expertise. When we compared the average dichotic reconstructions and classifications distributions of low and high musical expertise participants against the null distribution, we observed a difference for participants with high musical expertise but not for participants with low musical expertise. However, we found no main effect of musical expertise on both dichotic and monodic performances. Put together, these findings suggest that in the most challenging condition, i.e. the dichotic condition, musical expertise may have a facilitatory effect, but the effect may be small. This result partially corroborates previous works showing a better cortical tracking of music stimuli for musicians compared to non musicians (Di Liberto et al., 2020b; Doelling and Poeppel., 2015 ; Puschmann et al., 2019; Harding et al., 2019) nor the one showing better performances of musicians in dichotic listening (e.g. Barbosa-luis et al., 2021).

A possible reason for the small impact of musical expertise may rely on the importance of inhibiting irrelevant streams in our task. Indeed, a very recent study demonstrated that musical training was not associated with attention skills in a task where participants had to encode a target melody while inhibiting a distracting one (Blain et al., 2022, see also Bidelman and Yoo., 2020 for a similar conclusion with speech-in-noise task). While these results are in line with our findings, they should be nonetheless shadowed by the fact that this type of task also requires processes that are supposedly better in musicians such as stream segregation (e.g. François et al., 2014), cognitive control (e.g. Bialystok et al., 2009) or WM (e.g. Zuk et al., 2014). That being said, the superiority of musically trained people in executive functions is still under debate (e.g. Boebinger et al., 2015; Schellenberg., 2011 and Sachs et al., 2017 for evidence in children) and we found no clear differences between low and high musical experts in cognitive scores here. Another possible reason may rely on the choice of the stimuli. In the current setup participants heard simultaneously a different musical excerpt in each ear. However, the two excerpts were rather controlled in terms of rhythmic structure and spectral features (see Figure 1). Thus, while participants could rely on the spatial dimension to segregate the two streams, the spectral and temporal differences between the musical excerpts presented dichotically were small. By limiting the use of spectral and temporal cues in stream segregation, this may have reduced musicians’ advantage (Parbery-Clark et al., 2012; Fuller et al., 2014; Parbery-Clark et al., 2009a, b).

The main objective of this work was to explore whether executive functions, and more specifically WM, sustained attention and attentional inhibition, could explain the variability of reconstruction and classification of ecological monodic and dichotic stimuli based on neural data. We wanted to see whether the ability to sustain attention would impact the auditory tracking of the stimulus to attend. More precisely, participants with weak sustained attention abilities may experience more attentional disengagement periods (toward the stimulus to be attended), leading to poorer auditory cortical representation of the stimulus and therefore poorer AAD performance. However, our analysis revealed that sustained attention was not related to the performances in monodic and dichotic conditions. At least two non-exclusive explanations may be discussed here: the choice of the attentional task and the duration of the musical stimuli. First, as suggested by Helton (2009), SART performances, and more specifically errors of commission, can be seen as failures of motor control, while motor control may not be crucial when sustaining attention of auditory objects. Relatedly, compared to the continuous stream of music, the discrete stream of the SART may lead to differences in attentional maintenance and attentional capture (Yantis & Jonides, 1984; Sturm & Willmes, 2001; Esterman et al., 2013, 2019). Second, the common scores derived from the SART do not reflect moment-to-moment lapses of attention because they are mainly global measures of sustained attention over long periods of time. Thus, it could be that the SART was not able to capture the smaller scale sustained attention variations that were relevant when listening to the one-minute musical excerpts we used.

The WM task yielded similar results with both the monodic and the dichotic listening conditions. This result is less surprising. Indeed, it seems possible that in our listening paradigm where stimuli were long ecological polyphonic music excerpts, the ability to follow the stream of interest strongly relied upon attentional inhibition and to a lesser extent upon rehearsal, encoding or updating mechanisms. Although this seems evident for dichotic stimuli, one should recall that even in the monodic condition, the use of polyphony implies auditory stream segregation and integration (Wright & Bregman, 1987). Importantly, the high ambiguity of polyphonic music (Pressnitzer et al., 2011) is possibly mediated by an interaction of both bottom-up perceptual and top-down attentional processes (Bigand et al., 2000; Disbergen et al., 2018). While it has been proposed that a crucial function underlying the stable maintenance of items in WM is the ability to deal with distractors (Conway et al., 2001; Cowan et al., 2005; Bledowski et al., 2010), the WM task that we used (i.e. Backward Digit Span Task) does not measure this inhibitory component. This may explain why the WM score failed to predict the decoding performance in a task requiring complex auditory scene analysis. Clearly, this does not mean that WM is not involved in both monodic and dichotic listening (Peretz & Zatorre., 2005; Escobar et al., 2020 ; Colflesh & Conway, 2007; James et al., 2014; Janata et al., 2002). Further work is needed to more precisely understand how the different components of WM are involved in such complex situations such as listening to ecological polyphonic musical stimuli.

This interpretation seems to be comforted by the fact that attentional inhibition explained around 10% of the variance of dichotic reconstruction performance and around 8% of the classification one. This result confirms our hypothesis that a better ability to resist external distractors leads to better neural tracking of the auditory targets. This extends previous findings showing that cognitive control is involved in dichotic listening tasks (Hugdhal et al., 2009 ; Falkenberg et al., 2011) and auditory scene analysis (Goll et al., 2012). Moreover it seems to corroborate the manifold proposal about the implication of an inhibitory process in dichotic listening (see for example Conway et al., 2001 or Bledowski et al., 2010). The specificity of the link between auditory attentional inhibition and dichotic listening is also supported by the non-significant correlation between attentional inhibition score and performance during monodic listening, that is with no distractors. Overall, this result suggests that an attentional inhibitory mechanism is at work during dichotic listening and possibly affects the way the brain tracks the relevant auditory information. This extends to ecological music stimuli previous findings of auditory selective attention studies showing a decreased brain response of non-relevant simple sounds compared to relevant ones (Chait et al., 2010; Alain & Woods, 1994; Berman et al., 1989; Bidet-Caulet et al., 2007; Bidet-Caulet et al., 2010; Michie et al., 1990; Michie et al., 1993). In addition, this result is also in line with the aforementioned findings suggesting that maintaining a stable representation of the target stream during dichotic listening could be mainly based on the ability to deal with distractors. Nevertheless, further work has to be done to exactly characterize what type of inhibitory process is the most important during dichotic listening. Indeed, with the Stroop task we were only able to measure distractor resistance but it is not excluded that intrusion resistance could play a role in dichotic listening as well. Importantly, this is the first evidence that attentional inhibition, as assessed by a cognitive task, may have a predictive power of the performance of a neuro-steered hearing aid. This result also raises the question of whether training inhibition abilities may improve auditory neural tracking in complex auditory scenes.

Overall, the results of the presented study showed the feasibility of using the reconstruction stimulus approach with ecological polyphonic musical stimuli. Importantly, they point to specific cognitive mechanisms being at work during dichotic musical listening in normal hearing listeners. Further research should be carried on to investigate other cognitive functions that are involved in streaming (e.g. divided attention) as well as more precise sub-components of the executive functions, like distractor and intrusion resistance or updating and rehearsal in WM, in order to see to what extent they can predict the performance of the reconstruction stimulus approach. Also, it would be of interest to explore whether more general cognitive abilities such as general intelligence or cognitive flexibility could also be good predictors of AAD performance. In addition, It may be valuable to replicate this work with other populations such as CI implantees, children or the elderly, to see if the findings are consistent. Finally, as we show the predictability power of specific executive functions, it appears conceivable to study the impact of executive-function-based therapies to improve neuro-streered hearing aids performance.

In conclusion, one can say that cognitive factors seem to impact AAD performance and taking advantage of this relation could be useful to improve next-generation hearing aids. Thus, the combination of specific cognitive training and neuro-steered hearing aids seems to be a serious option to help deaf people perceive the world a little more naturally.

## Acknowledgements

We wish to thank Sophie Chen and Christelle Zielinski for technical help in synchronizing auditory triggers and EEG signals, Aurélie Bidet-Caulet for very helpful comments on a previous version of this manuscript and Samuel Deslauriers-Gauthier for mathematical help in the creation of the Dichotic Performance Index.

J.B. is funded by Oticon Medical (research agreement n°14294-CRI04) and Region Provence-Alpes Côte-d’Azur. D.S. was supported by APA foundation (RD-2016-9), ANR-16-CE28-0012-01 (RALP), ANR-CONV-0002 (ILCB), ANR-11-LABX-0036 (BLRI) and the Excellence Initiative of Aix-Marseille University (A*MIDEX).

